# Terpene volatiles mediates the chemical basis of blueberry aroma and consumer acceptability

**DOI:** 10.1101/2021.12.24.474137

**Authors:** Luis Felipe V. Ferrão, Haley Sater, Paul Lyrene, Rodrigo R. Amadeu, Charlie Sims, Denise Tieman, Patricio R. Munoz

## Abstract

Among the main features treasured by blueberry consumers, flavor is the most important. Human perception of food flavors can primarily be divided into two main sensory inputs, taste and aroma. Through retronasal olfaction, a group of metabolites called volatile organic compounds (VOCs) emitted from the fruit are able to produce the sensation of aroma, creating the myriad of flavors experienced during our life. In blueberry, breeders have noticed some genotypes with unique floral and sweet flavor notes that, ultimately, enhance human aroma perception. Despite the importance, both the understanding of which chemicals are mediating this variation across phenotypes and the potential impact on consumer preferences remains largely unknown. In this study we dissected the main components underlying blueberry aroma and associated it with consumer predilections by paring metabolomics with sensory analysis. Our contribution in this study is four-fold: (i) first, based on a representative blueberry germplasm cultivated at the University of Florida, we differentiated genotypes with floral and sweet aromatic notes and confirmed that such unique characteristics are preferred by consumers; (ii) at the chemical level, we showed that a group of eight terpene volatiles constitute the primary metabolic group associated with aroma sensation; (iii) we demonstrated that aromatic genotypes can be classified using information from a group of a few key volatiles; and finally, (iv) we combined pedigree and metabolomic information in a single predictive framework and showed the importance of metabolomic data for flavor-assisted selection. For the blueberry community, our findings open new venues to explore flavor. Broadly, we present an emerging view about flavor and provide a detailed blueprint of how this target could be addressed in fruits and vegetables.

## 1 Introduction

Fruits are valuable sources of beneficial bioactive compounds in the human diet and are an integral part of healthy lifestyles. Over the last decades, a flurry of publications have discussed the psychological factors that modulate our personal predilections for food consumption (Kader, 2008; Bruhn *et al*., 1991; Klee, 2010; Klee and Tieman, 2018). Better-tasting foods are arguably among the most important attributes, making the flavor phenotype an important target for breeding. Despite the significance, headway in this field is not simple and has experienced slow progress in recent decades due to the complexity of measuring the “flavor phenotype” (Klee, 2010). Human perception of flavor is subjective and depends on the integration of multiple sensory systems, specifically taste and olfaction, which can create a myriad of flavor experiences (Klee and Tieman, 2018; Goff and Klee, 2006; Schwab *et al*., 2008). In this scenario, the challenge faced by modern breeding programs is to disentangle the chemical compounds that effectively contribute to human liking and, more importantly, use this information to select and release cultivars with improved sensory perception by consumers.

Blueberry (*Vaccinium* spp.) is among the most consumed soft fruits and has been recognized as an important source of health-promoting compounds. Numerous reports have shown that the species is a rich source of bioactive and antioxidant compounds, which mostly support blueberries’ current status as a global ‘superfood’ (Kalt *et al*., 2019; Norberto *et al*., 2013). A previous psychophysical survey suggested that blueberry consumers consider flavor as the most important attribute for fresh consumption (Gilbert *et al*., 2014). With a relatively recent domestication history, flavor is a trait that still possesses wide genotypic variability for exploration in the *Vaccinium* genus. More specifically, a large diversity of pleasant flavor notes – that include fresh green, fruity, floral and tea-like aromas – have been systematically observed within the blueberry germplasm and classified as ‘aromatic genotypes’ – a term borrowed from the breeding jargon. Although remarkable, the real impact of these unique aroma notes remained elusive, mostly due to the absence of evidence whether these aroma notes could be perceived (and appreciated) at the consumer level. To fully grasp how these aroma notes might affect human preferences, we first need to understand the link between the flavor chemicals of a fruit and hedonic ratings from consumers.

Among the main factors driving flavor perception in blueberry fruit; sugars, acids and volatile organic compounds (VOCs) are the most important (Klee and Tieman, 2018; Gilbert *et al*., 2015). Sugars and acids are detected by taste bud receptors on the tongue which results in the human sensation of taste. In contrast, volatiles are perceived by our olfactory system creating the sensation of aroma(Klee, 2010; El Hadi *et al*., 2013). It has been shown that aroma contributes much of the diversity of flavors we experience, thereby leveraging the importance of VOCs to breeding (Klee, 2010; Goff and Klee, 2006). The delicate aroma of ripe fruits is formed by a unique combination of hundreds of VOCs derived from primary and secondary metabolites. In blueberries, many surveys have been dedicated to explore this volatile profile and many hundreds of VOCs from different pathways have been reported over the years (Sater *et al*., 2020) (Farneti *et al*., 2020). However, only a few of these studies have sought to compare human evaluations with the volatile profile of blueberries (Gilbert *et al*., 2015; Du *et al*., 2011) and so far there is no consensus as to which VOCs are both impacting blueberry flavor and influencing consumer liking. These studies not only reinforce that flavor is complex, but also show the diversity of compounds that can be harnessed to enhance and even improve the flavor profile of blueberries through breeding.

Aiming to advance our understanding of human flavor preferences for blueberry, we paired sensory and metabolomic analysis using a panel of genotypes representative of the active germplasm planted around the world and maintained at the University of Florida (UF) breeding program. Starting in 1950, UF has greatly contributed to expanding production areas by developing low-chilling cultivars. So-called southern highbush blueberry, this group of cultivars is better adapted to subtropical and Mediterranean climates, and represents almost 40% of all the blueberry globally produced currently. Given its importance and motivated by observational evidences that some genotypes consistently presented unique floral and sweet aromatic notes, we addressed the following specific questions in this study: (i) do consumers perceive and prefer berries with such unique aromatic notes?, (ii) what are the main chemical compounds that dictate such aromatic sensation?, (iii) from the breeding standpoint, is aroma preference predictable? and, more broadly, (iv) can flavor preferences be better predicted by combining pedigree and metabolomic information? We anticipated that consumers would rate aromatic berries with higher liking scores, while metabolic profiling indicated that a group of terpene volatiles were positively associated to this flavor perception. Importantly, the methods and approaches we used here allow us to provide a blueprint of how blueberry flavor can be understood and opens the frontier to using statistics to predict human flavor interpretation using only a few metabolites.

## 2 Material and Methods

### 2.1 Plant Material

In this study, we used 52 Southern Highbush Blueberry genotypes from the University of Florida (UF) breeding collection. These consisted of advanced selections, several cultivated genotypes and interspecific hybrids cultivated in two representative locations of the Floridian blueberry production: the Plant Science Research and Education Unit in Citra (CI) (latitude 29024’46.0” north, longitude 82006’35.5” west) and Waldo (latitude 29047’23.7” north, 82010’04.9” west). In 2017, to select genotypes that presented unique flavor notes, an internal panel composed by qualified blueberry breeders scored aroma using a 0-5 scale (where 0 represented ‘no aroma intensity’ and 5 ‘extreme aroma intensity’) during the season. This first screening effort was used to establish a group of 21 highly flavorful genotypes that, among other factors, presented distinguishable floral and sweet aroma notes (Bett-Garber and Lea, 2013). As a reference, breeders seek aromatic notes that resemble the aroma of the cultivar “Kestrel” – a blueberry cultivar released by the UF breeding program that has been consistently rated by specialists and consumers with high scores of overall liking and flavor intensity. Thus, we borrowed a term from the breeding jargon and classified 21 genotypes as “aromatic”, while the remaining 31 blueberry genotypes which lacked these distinctive aromas were considered “non-aromatic”.

In 2018 and 2019, fruit was harvested twice per week when fully ripe, beginning the first week of April and continued until May 8th and May 4th, respectively. From each individual adult plant, berries were harvested in bulk to make a representative fruit sample then subdivided into batches designated for sensory panels and biochemical measurements. All berries were stored overnight at 4^0^C before being sent for sensory panel tasting or for fruit quality analysis and volatile quantitation.

### 2.2 Sensory analysis

Over the course of two years, 52 genotypes were subjected to sensory evaluation in 16 different panels, with an average of 84 panelists per sensory panel. Consumers panels were conducted at the Food Science and Human Nutrition Department at UF, Gainesville, FL. As previously described by Gilbert *et al*. (2015), each panelist scored five to six genotypes in a single panel arranged in a Williams design and generated by Compusense 5.6 (Compusense Inc., Guelph, Ontario, Canada). Panelists were given cups with three to five berries of each genotype to taste and asked to evaluate their (i) liking preference for each berry sample, as well as the (ii) intensity of experience. The Global Labeled Magnitude scale (GLMs) (0 to +100; 0 = no sensation, 100 = most intense sensation of any kind) and the Hedonic Global Labeled Magnitude scale (hGLMs) were used (Bartoshuk *et al*., 2004, 2005). By using the hGLM scale the panelist evaluated the overall liking, flavor liking, and texture liking; while intensity of sweetness, sourness, flavor, firmness and juiciness of each genotype were scored using the GLM scale.

Additionally, the panelists were asked simply to determine whether they could discern aromatic and non-aromatic genotypes using the following questions: (i) *“Some blueberries have unique or stronger flavor profiles than others. Do you detect any fruity, floral or other novel flavors in this blueberry?”*; and (ii) *“If you answered yes to the last question, please describe on the lines below what qualities of the berry flavor was unique. Example: This berry sample had a floral note, or this berry sample had abnormally strong blueberry flavor*.*”* Both questions were used as a qualitative measure to understand, if prompted, whether the panelist could differentiate the breeder predetermined classification between aromatic and non-aromatic genotypes. In order to accommodate the consumer classification of unique flavor from the first question, we used panelist’s binary (yes or no) response to establish a new trait referred to as ‘Novel Flavor’

### 2.3 Biochemical Analysis

Volatile collection was performed in a larger sample of genotypes than was included in the sensory analysis. A total of 70 genotypes were used in the biochemical analysis including the 52 genotypes used in the sensory analysis. Volatiles were extracted using 100 grams of washed and dried ripe berry tissue. Berries were sliced in half to allow fruit flesh to exude volatiles. Using a method developed by Schmelz *et al*. (2003) and adapted by Tieman *et al*. (2006), sliced berries were subjected to dynamic headspace trapping for 1h. Volatiles were trapped on a SuperQ column and eluted using 150 *µ*l methylene chloride (75-09-2), and 5 nonyl acetate (143-13-5) as an internal standard for elution normalizing. Samples were frozen at -80 C until volatiles were separated using a DB-5 column on a Agilent 6890N gas chromatograph attached to an Agilent 5975 mass spectrometer (Agilent, Palo Alto, CA). Volatile quantitation was preformed using ChemStation software (Agilent, Palo Alto, CA). Volatile data from the GC-MS were translated using GC Method Translation Software (Agilent, Palo Alto, CA). Files from each sample were quantified using MassHunter Software 10.0 (Agilent, Palo Alto, CA). Compounds were initially identified through NIST 11. Identification and quantification of each compound was performed using a calibration curve made from analytical standards. Effectively, compound concentration was calculated as the signal response for each peak relative to the concentration curve for that compound standard and normalized by the signal response of the internal standard. Nonyl acetate was used as the internal standard for all compounds consistent with the methodology used by Gilbert *et al*. (2015). Volatile concentrations are in units of ug g-1 FW h-1. Multiple technical replicates per sample were analyzed by GC-MS, and average quantification values per genotype within the season were used as phenotype. Five non-volatile fruit quality traits were also measured using the protocols described by Cellon *et al*. (2018) AND FerrÃo *et al*. (2018), including soluble solids, pH, titratable acid (TA), size and compression force.

### 2.4 Statistical Analysis

#### 2.4.1 Sensory and biochemical data analysis

At the sensory level, hedonic traits evaluated in the 52 genotypes were analyzed using information from both seasons (2018 and 2019). A univariate linear mixed model with season as a fixed effect, and panelist nested in panel session as a normally and independent random effect, was used. To estimate variance components the genotype effect was firstly treated as a random effect with variance proportional to a pedigree relationship matrix **A**. The pedigree matrix was computed as described by Cellon *et al*. (2018). Outliers were removed if the absolute value of standardized residuals was larger than two residual standard deviations. To estimate adjusted means for all subsequent analyses, the genotype effect was considered as a fixed effect. Mixed models were implemented using the `asreml-R` 4.0 software (Butler *et al*., 2009), while the **A** matrix was computed using the `AGHmatrix` software (Amadeu *et al*., 2016).

Differences between aromatic and non-aromatic phenotypes per sensory traits were quantified via analysis of variance (ANOVA). To this end, the adjusted means estimated per genotype and pre-corrected for season and panelist effect were used as the response variable, while the sugar and acid ratio, berry size and compression force (proxy of firmness) were used as cofactors. Analyses were performed in R using the `lm()` function. Additionally, a multivariate analysis was carried out for the description data provided by the panelists. For that, we encourage the panelist to make comments on remarkable samples during the sensory panels. We processed comments by regrouping words and summarizing the most common terms used to describe genotypes in the aromatic and non-aromatic group.

A similar strategy for the sensory data was used to estimate variance components and the adjusted means for the 70 genotypes for which volatiles were collected. To this end, a univariate linear mixed model was fitted accounting for the fixed effect of season and the genotypic effect. All volatile data were previously log transformed. After that, the adjusted means estimated for the 52 genotypes that are in common in both sensory and chemical were used for Principal Component Analysis (PCA), as implemented `FactoMineR` package (LÊ *et al*., 2008). Differences in sensorial and biochemical variation between the aromatic and non-aromatic groups were accessed via ANOVA models using the lm() R function.

### 2.4.2 Inference and Prediction

Statistical inference and estimation were performed by combining Ridge Regression and Random Forest regression models, where we related sensory attributes (response) to the biochemical data (predictor). A similar approach was originally described by Colantonio *et al*. (2020). Thus, for the Ridge Regression analysis, we determined the direction (negative or positive effect) of each biochemical component on sensory variation. A cross-validation scheme was used to optimize the regularization parameter in the Ridge Regression using the `MASS` package (Venables and Ripley, 2013). Concurrently, Random Forest was used to estimate the importance of each individual metabolite. Random Forest regression models were implemented using the default options implemented in the `randomForest` package (Breiman, 1996) (number of trees to grow equal to 500 and number of variables randomly sampled equal to p/3, where p is the number of variables), followed by a permutation test, as suggested by Altmann *et al*. (2010) and implemented in the `Vita` R package.

To investigate variance components, we relied on a multi-kernel mixed model, where we assume the outcome (sensory evaluation) can be modeled as a sum of genetic and chemical’s effects. For this, we considered three main sources of information: pedigree (accessed via kinship records), instrumental data (including only the non-volatile data; soluble solids, TA, pH, size and compression force) and volatile data. To access the importance of these source of information in predicting sensory evaluations, we first estimated variance components using a suite of kernel functions. To this end, the following linear mixed model was used:

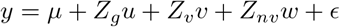

where *y* is the response variable and represent pre-corrected sensory evaluation, 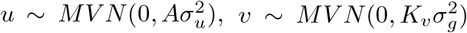, and 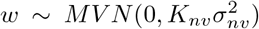 are independent random effects, normally distributed and with variance and covariance terms defined as kernel matrices constructed using pedigree, volatile and instrumental covariables, respectively. We use *Z*_*g*_,*Z*_*v*_ and *Z*_*nv*_ to denote the design matrices associated to the genetic, volatile, and non-volatile data. For the genetic information, the **A** pedigree relationship matrix was computed as previously described. For volatile and instrumental traits, the *K*_*v*_ and *K*_*nv*_ matrices are Gaussian kernels defined in the Euclidean space and constructed as described by Colantonio *et al*. (2020). Residual term was assumed to be normally and independently distributed 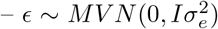. The terms 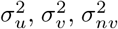, and 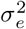 are representing the variance components associated to the pedigree, volatile, non-volatile and residual random effects, respectively.

For prediction analyses, we used a linear mixed model relied on genomic best linear unbiased prediction (GBLUP) (Meuwissen *et al*., 2001), as previously suggested by Longin *et al*. (2020). Therefore, we assumed the following mixed linear model:

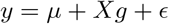

where *y* is the response variable and represent pre-corrected sensory evaluation, **X** is the incidence matrix linking observation in the vector *y* to their respective genotype effects in the vector g. Normality was assumed for the multi-omics and residual effects, where 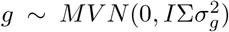 and the residual 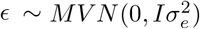. For the multi-omics term, the Σ variance and covariance structure was defined by combining different proportions of pedigree and biochemical (volatile and non-volatile) data. For that, kernel computed using the pedigree and biochemical information (volatile and non-volatile data) were blended into a single kernel matrix, representing the relationship among genotypes as given by these two sources of information. To this end, the following equation was used: Σ = *wA* + (1 − *w*)*K*, where **A** represents the pedigree relationship matrix and **K** is a Gaussian kernel defined in the Euclidean space and computed using the biochemical (volatile and non-volatile) data. The percentage of pedigree data used to create this new kernel is defined by *w* in the equation and it ranged from 100% (representing a traditional **A** matrix) to 0% (kernel matrix based solely on the biochemical data). Predictive abilities were accessed via a 10-fold cross-validation scheme and computed using the Pearson’ correlation between the adjusted means computed for the sensory traits and the predicted values. Predictive analyses were implemented using the `asreml-R` 4.0 software (Butler *et al*., 2009).

## 3 Results

To evaluate the metabolite and sensory differences between the 52 blueberry genotypes classified by breeders as either “aromatic” or “non-aromatic”, we accessed a total of 68 volatiles and connected them to eight sensory evaluations (Table S1 and Supplemental Figure S2). Among the sensory traits, we first observed that genotypes classified as “aromatic” were preferred by blueberry consumers when compared to “non-aromatic” fruit. By accessing differences in the biochemical profile between the two groups, we quantified eight differentially expressed terpene volatiles that potentially play a key role in blueberry aroma. Finally, predictive performances of aroma and other sensory traits were posteriorly accessed using linear mixed models combining genetic and metabolomic data. While pedigree information was less effective at forecasting sensory perceptions, the volatile information showed good prospects to be used in metabolomic selection for flavor improvements.

### 3.1 Blueberry genotypes vary in sensory scores and biochemical profiling

Volatile profiles were accessed in a group of 70 genotypes maintained at the UF breeding program, where 52 were subjected to taste panels in 2018 and 2019. Among the tested genotypes, the selection FL10-144 (aromatic) presented the largest adjusted mean for overall liking and sweetness intensity (Table S2). Another aromatic genotype, FL06-510, presented the highest adjusted means for both flavor intensity and flavor liking. In contrast, “Vireo” (a non-aromatic genotype) had the lowest scores for both overall liking and flavor liking, and the highest sourness perception (Table S2). Importantly, for all sensory traits we observed large variability, but with low associated values of narrow sense heritability (Table S2).

In total, 68 volatiles were identified and quantified via dynamic headspace gas chromatography-mass spectrometry (GC-MS). From this initial set, 40 VOCs (59%) remained after the removal of metabolites that did not show variability in this population. The remaining VOC compounds were classified based on their biosynthetic origins, including 13 terpenes, nine oxylipins, nine amino acid derived compounds, four fatty acid derived, three phenylpropanoids, two methylketones, and two of unclassified origin – likely from primary metabolism (Figure 1a). The most abundant class observed in both ‘aromatic’ and ‘non-aromatic’ groups were oxylipins and methylketones (Figure 1b). Remarkably, the relative concentration of terpenoid compounds in the “aromatic” group was four times larger than in the “non-aromatic” berries (Figure 1b). In total, eight volatiles from the terpenoid group showed concentrations statistically different across the “aromatic” and “non-aromatic” groups: P-cymene (PCYM), myrtenal (MYRTENAL), linalool (LINALOOL), (L)-carvenol (LCARV), geranyl acetone (GEROTE), geranyl acetate (GERATE), D-limonene (DLIMON) and beta-myrcene (BETAMYR) (Figure 1a). A lexicon of sensory descriptors associated to the blueberry volatile profile are organized in the Supplemental Figure S1.

**Figure 1:**
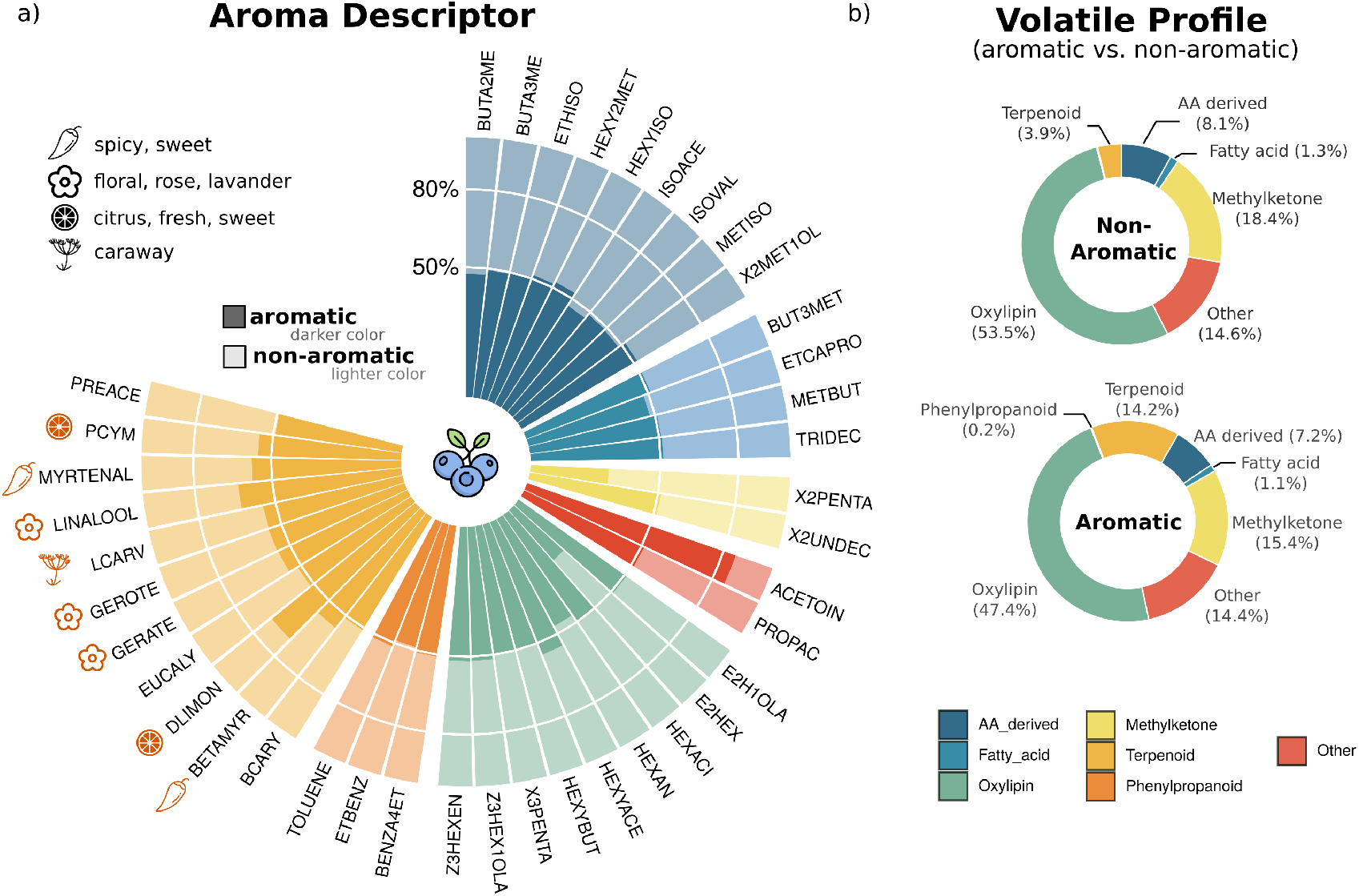
a) Relative concentration of each volatile. Values per volatile were first normalized to range from 0 to 100% and projected to show differences across “aromatic” and “non-aromatic” groups. Analysis of Variances were performed per volatile, and significance differences (P-value ≤0.1) across the ‘aromatic’ and ‘non-aromatic’ groups were observed for the P-cymene (PCYM), myrtenal (MYRTENAL), linalool (LINALOOL), (L)-Carvenol (LCARV), geranyl acetone (GEROTE), geranyl acetate (GERATE), D-limonene (DLIMON) and beta-myrcene (BETAMYR) volatiles. More details on the odor descriptors of the eight terpene volatiles is presented in the Figure S1 and the full name of the volatiles are presented in the Table S1. b) Average concentration per chemical group within the “aromatic” and “non-aromatic” group are classified by their relative proportions for each biosynthetic metabolic group.

Volatile distributions appear to vary significantly between genotypes and even within a single biosynthetically related group of compounds. Medium-to-high narrow sense heritability values were observed for most of the metabolites (Table S3). An average heritability value of 0.30 was observed across all volatiles. The largest value was observed for the ISOBISO (isobutyl isobutyrate) (0.91), a branched chain amino acid volatile. Among the terpenoids that showed a statistical difference between both groups, the heritability value associated to P-cymene (PCYM) showed the largest value (0.59), while geranyl acetone (GEROTE) had the lowest value (0.12).

### 3.2 Sensory analysis indicates that blueberry consumers prefer aromatic berries

Our first research question addressed whether aromatic berries could be perceived by consumers. Overall liking for the “aromatic” group was higher and statistically different than the “non-aromatic” group (P-value ≤ .05) confirming our initial hypothesis that such unique flavor notes are perceived differently by consumers and have a positive flavor sensation (Figure 2a and Table S4). Four of the other sensory traits examined – including flavor liking, sweetness intensity, juiciness intensity, and flavor intensity – had higher average scores for aromatic berries as opposed to non-aromatic berries. Only sourness exhibited similar distribution patterns in both groups. Despite the difference in sweetness intensity, “aromatic” and “non-aromatic” genotypes, on average, did not present significant differences (P-value ≥.05) in soluble solid content. This ultimately suggests that other chemical attributes are enhancing the sweetness perception at the sensorial level. As a complement, by using a word cloud analysis, we accessed patterns in the panelist’s comments regarding their sensory experience with the different genotypes (Figure 2b). Words referring to positive experience and flavor notes were used more frequently for “aromatic” genotypes, including the terms “floral”, “flavor” and “fruity”. Notably, words used to express negative flavors (or experiences) were more frequently used to describe genotypes within the “non-aromatic” group, including “weak”, “watery”, “bland” and “bad flavor”.

**Figure 2:**
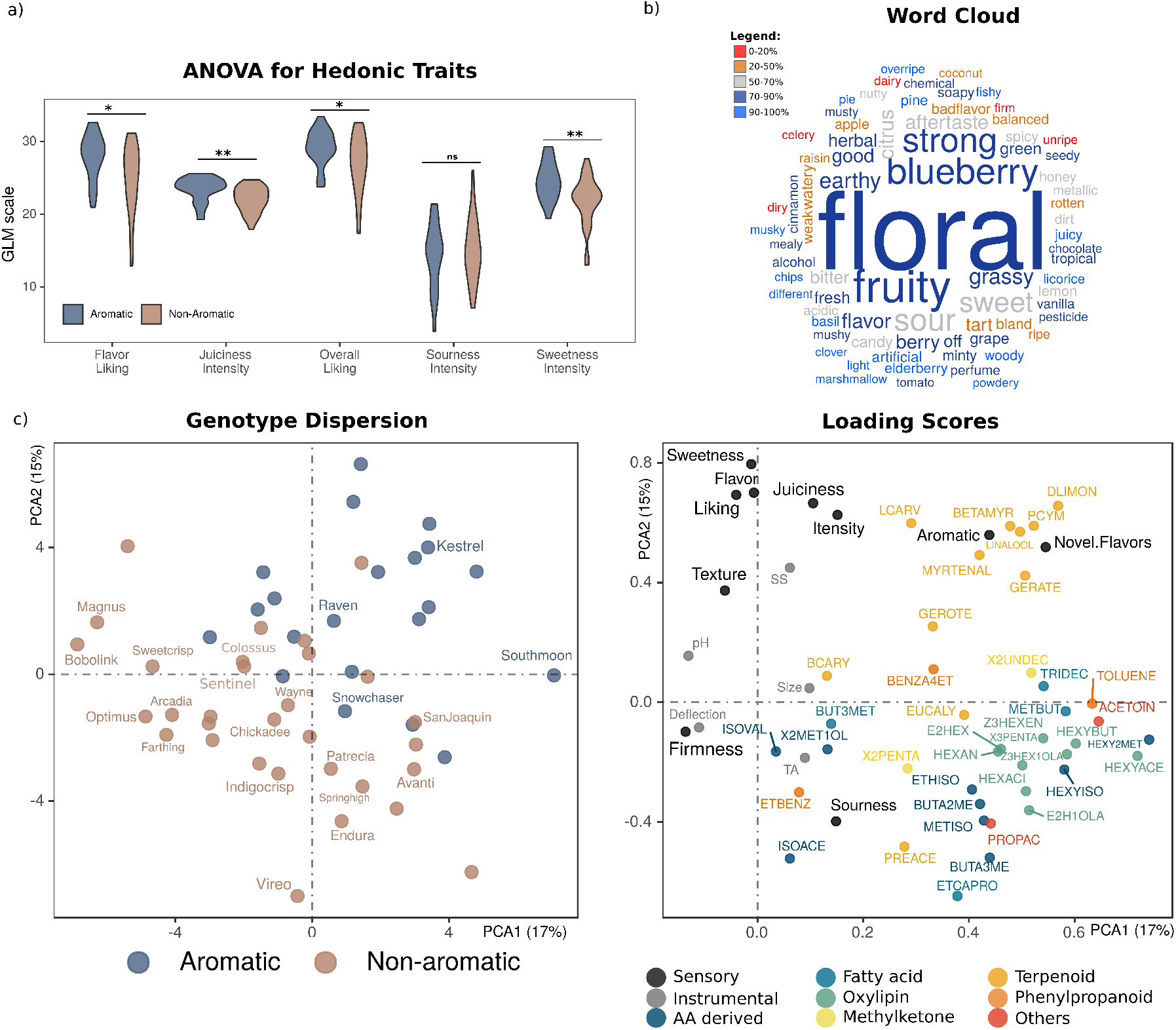
a) Analysis of Variance (ANOVA) contrasting mean sensory traits evaluated in the aromatic and non-Aromatic groups, where **, *and ns refer to P-values ≤.01, P-values*<*.05 and P-values ≥.05. b) Panelists comments represented via word clouds where size of the word was proportional to the frequency that a given word was used and color represents weather each word was more used to describe aromatic genotypes (blue) or non-aromatic (red). c) Principal Component analysis representing the genotype classification (left) and the loading scores (right) – blueberry cultivars released by the University of Florida are highlighted in the genotype classification plot, while the other dots are representing advanced selections in the blueberry breeding germplasm.

Principle component analysis (PCA) was conducted to group the genotypes by integrating sensory, volatile and non-volatile information (Figure 2c). By using the adjusted means of 52 genotypes subjected to sensory and biochemical analysis, we observed that most of the aromatic berries formed a unique group residing in the first and second quadrants of the PCA plane. The PC loading scores showed the importance of each sensory and chemical component. Overall liking is positively correlated to soluble solids and sensory traits, including sweetness, flavor and juiciness. As might be expected, firmness perception and compression force were also positively correlated. Texture grouped independently, which is expected given the difficulty for untrained panelists in describing fruit texture.

Importantly, a group of primarily “aromatic” genotypes towards the first quadrant of the PCA correlated to the ‘Aromatic’ and ‘Novel.Flavor’ sensory traits. ‘Aromatic’ stands for the breeder classification for unique flavor notes, while analogously ‘Novel Flavor’ refers to the binary trait scored by consumers. Both traits are highly correlated suggesting agreement between panelists and breeders. At the metabolite level, both response variables are positively correlated with terpene volatiles, including LINALOOL, DLIMON, BETAMYR, MYRTENAL, PCYM, GEROTE, GERATE and LCARV. Many of the terpenes have similar odor descriptions as those identified by consumers in the word cloud (Figure 2b), including “floral” and “green” notes (Figure 1a). In contrast, “non-aromatic” genotypes were primarily distributed in the bottom two quadrants of the PCA.

### 3.3 Terpenes play an important role on enhancing aromatic perception

By using a combination of linear and non-linear regression models, we quantified the importance of chemical components related to human sensory perception. To this end, we first interpreted the slope coefficients estimated via ridge regression to indicate the direction (positive or negative) and Random Forest to regulate the importance of each compound (Figure 3a and Supplemental Figure S3). Instrumental traits including soluble solids, TA and PH, were important to explain variation in overall liking, sweetness and sourness intensities. On the other hand, size and compression force were more important variables to explain physical attributes associated to texture and firmness intensities (Supplemental Figure S3). Volatiles were more important to explain sweetness and juiciness intensities; and, especially, aroma sensations. As evidenced in the PCA analysis, a group of several terpenes are positively associated to the aromatic classifications (‘Novel Flavors’ and ‘Aromatic’ sensory descriptors defined by consumers and breeders, respectively).

**Figure 3:**
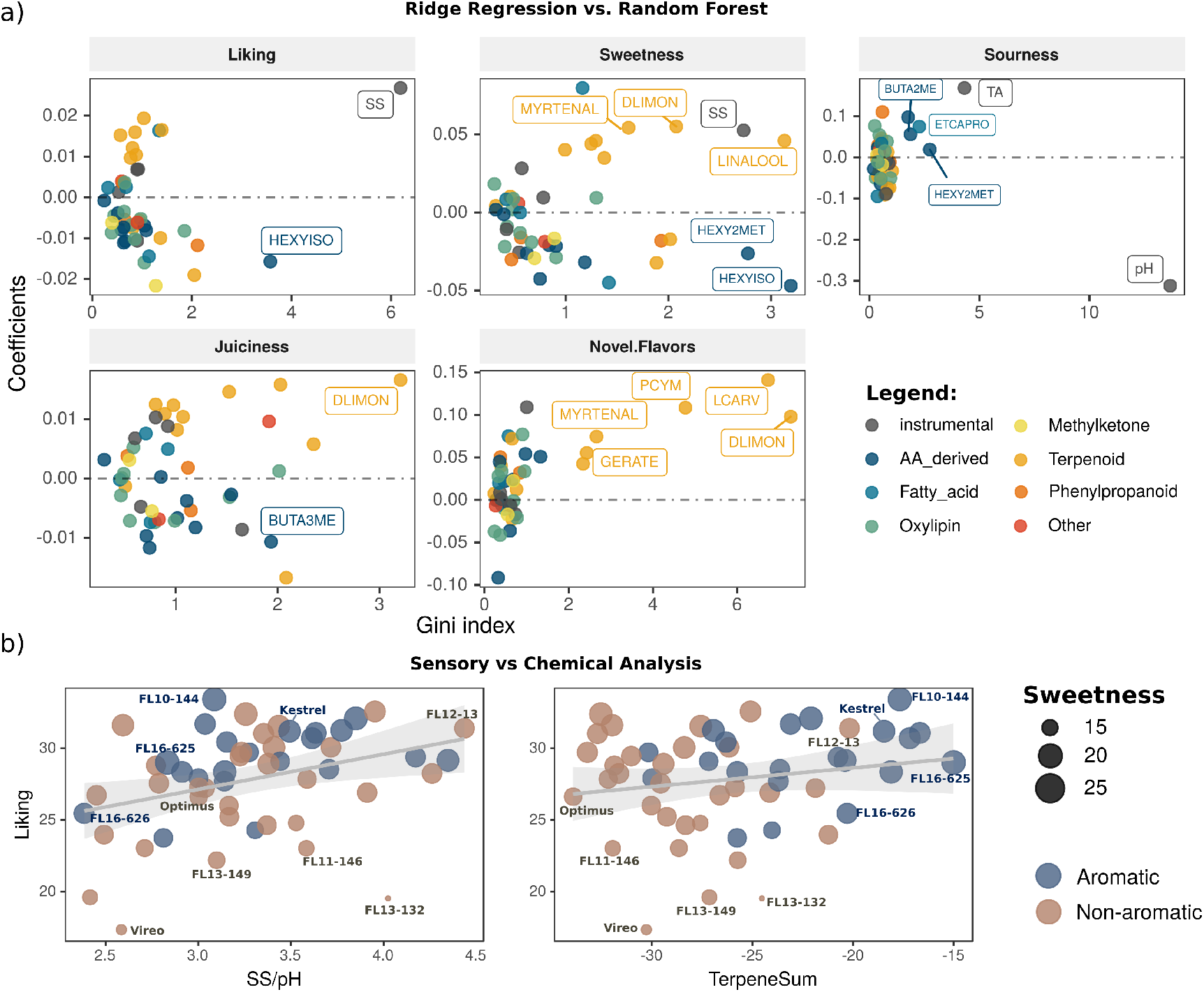
a) Regression coefficients estimated using Ridge Regression and variable importance represented by the Gini Index estimated via Random Forest regression models. b) Scatter plots relating the adjusted means for overall liking and SS/pH ratio and total terpenes (presented in the log10 scale). Size of the circles stands for the adjusted means estimated for sweetness.

Recently, it was proposed that the blueberry sugar and acid ratio can be used as a predictor of blueberry liking (Gilbert *et al*., 2015). We expanded such evidence by generating an integrated scatter plot where overall liking was regressed as a function of the sugar and acid ratio (SS/pH) and total terpenes (Figure 3b). The positive and linear trend observed for the sugar and acid ratio is expected and confirms preliminary expected. However, the positive relationship between terpene concentration and overall liking sheds some extra light on the relevance of flavor-related compounds that are not sugars or acids. To bring focus on the relative importance of terpene volatiles, we highlight some genotypes. For example, FL10-144 and FL13-149 are both genotypes with similar sugar and acid ratios, but different liking and sweetness scores. In counterpart, FL16-625 is a genotype with large sweetness perception, but low-to-moderate sugar content. Thus, we hypothesized that terpene volatiles make contributions to perceived sweetness independent of sugar concentration. However, not all terpenes are positively enhancing sensory sensations: eucalyptol (EUCALY) and prenyl acetate (PREACE) are some terpene volatiles that negatively impacted overall liking (Figure 3a and Supplemental Figure S3).

### 3.4 Aroma perception can be predicted using a few groups of terpene volatiles

With the understanding that terpene volatiles are modulating consumer acceptability, we evaluated an additional classification step by using a few groups of terpene as the input to classify genotypes as “aromatic” or “non-aromatic” and therefore resemble the breeder’s classification. Using logistic regression models and receiver operator characteristic (ROC), we computed the area under the curve (AUC) for seven terpenes and determined the optimal cutoff values for predicting aromatic genotypes (Figure 4). Myrtenal (MYRTENAL), P-cymene (PCYM) and D-limonene (DLIMON) presented the highest AUC values and hence are our best candidates for classification using few genotypes. Our empirical results suggested that, by ignoring eventual interaction effects that might mask the aromatic trait, genotypes can be classified as “aromatic” with a minimum of 80% of sensibility (true positive rate) and 70% of specificity (false negative rates) using only three volatiles. Thus, using this model, a genotype was classified as aromatic when the concentration of MYRTENAL, PCYM and DLIMON was larger than 0.13 ug g-1 FW h-1, 0.17 ug g-1 FW h-1 and 0.08 ug g-1 FW h-1, respectively. As a crude comparison, we ran the same classification exercise via logistic regression, but using other chemical components. Poor classifications were visually achieved when instrumental parameters (including sugar and acids attributes) or even non-terpene volatiles were use in the ROC analyses (Figure 4).

**Figure 4:**
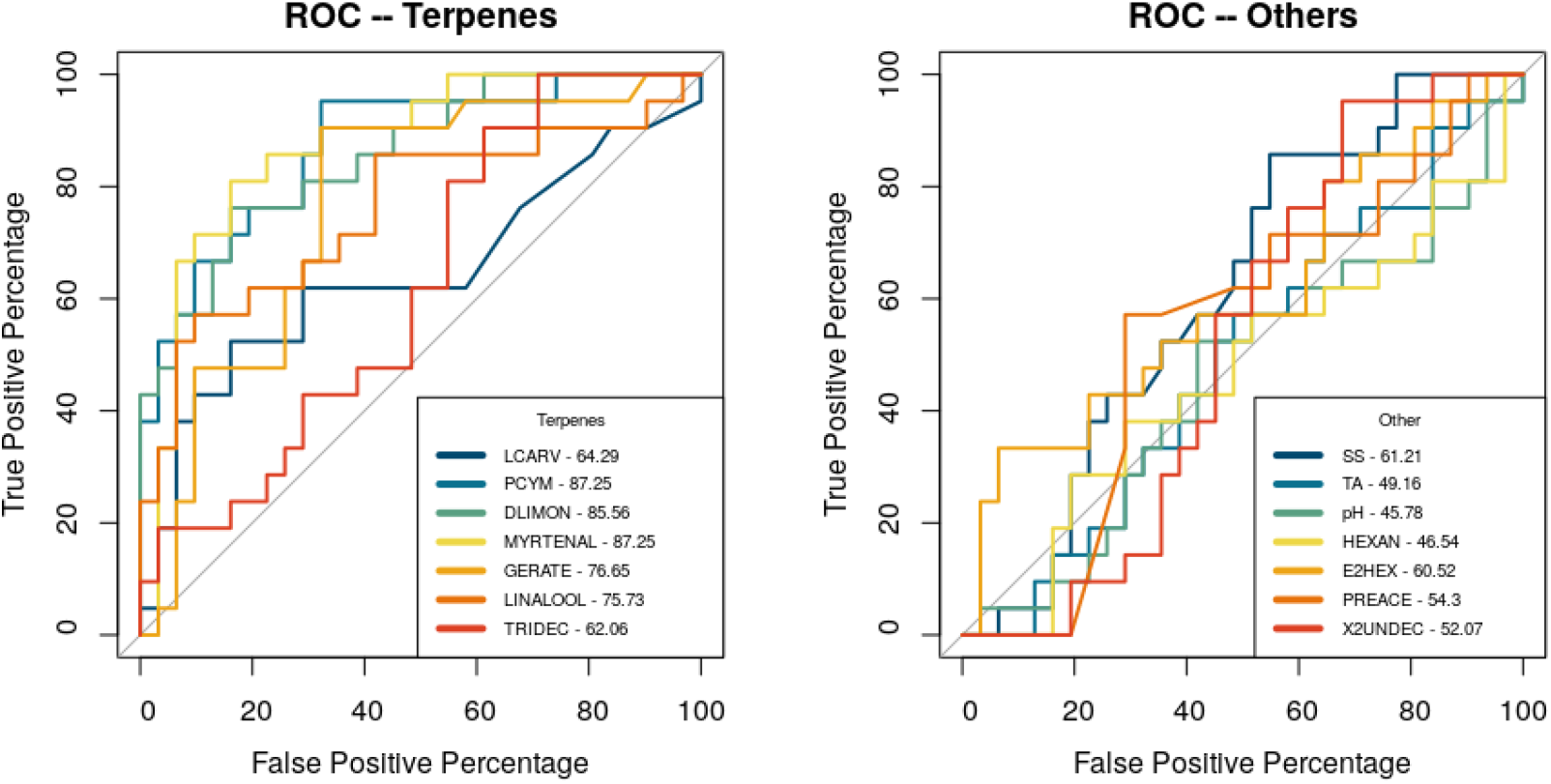
Comparison of classification performance for blueberry aroma. Classification was carried out using logistic regression models and receiver operator characteristic (ROC), while the performance is measured by area under the curve (AUC), where a higher value indicates better performance in classifying blueberry genotypes as “aromatic”. In the left plot, AUC is reported for seven terpene volatiles associated to the aromatic blueberry phenotype. Comparatively, the AUC values are also presented for some instrumental traits (SS, TA and pH) and non-terpene volatiles indicating poor classification, when contrasted to the AUC results for the terpene volatiles (right). Chemical name and AUC values are presented in the legend.

### 3.5 Flavor traits accessed via consumer panels are more accurately predicted using metabolomics data

After measuring the importance of metabolites on aroma acceptability, we assessed the importance of pedigree, volatile and non-volatile information to explain the variation of other sensory traits associated to flavor (Figure 5a). To this end, we first considered all volatiles and non-volatiles simultaneously on RKHS models. The additive genetic component accounted for the pedigree information did not explain large portions of the phenotypic variation observed in sensory characteristics. These results are consistent with the low heritability values previously reported for sensory traits (Table S3). Physical sensory attributes related to firmness and texture intensities had the largest impact in prediction accuracy by incorporation of pedigree information. In sharp contrast, non-volatile data were important sources of information to explain variation such as texture, firmness, sourness, and liking sensory traits. Volatiles played an important role to explain variation in aromatic sensation (‘Novel Flavor’) and sweetness intensity.

**Figure 5:**
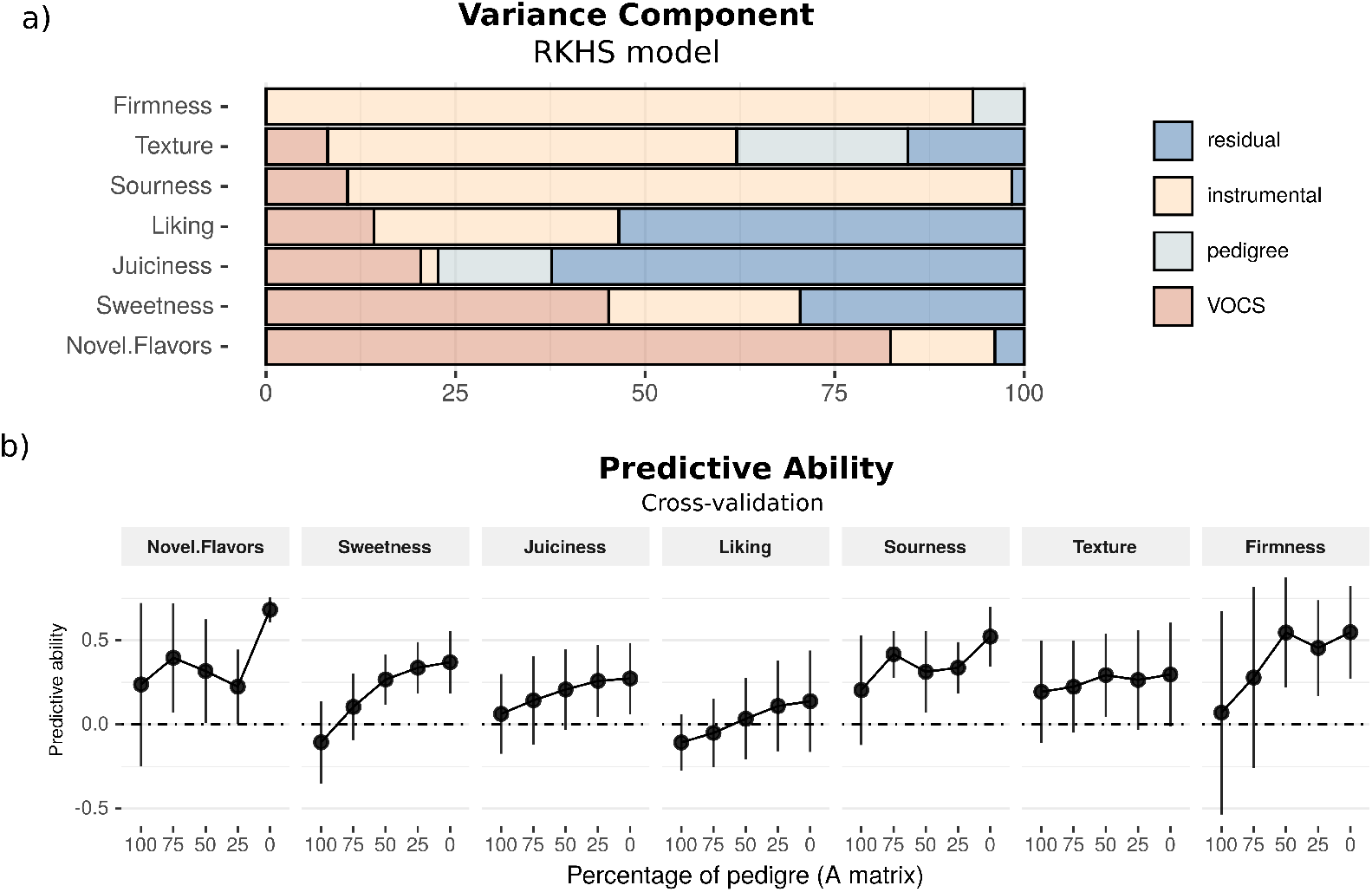
a) Variance component decomposition for seven sensory traits evaluated in blueberry using reproducing kernel Hilbert spaces (RKHS model) with kernels constructed using the pedigree, instrumental (soluble solids, TA, pH, compression force and size) and volatile (VOCs) data set. b) Predictive ability accessed via 10-fold cross-validation using pedigree and metabolomics information. Values in the x-axis stands for the percentage of pedigree information used to compute the variance and covariance matrix in the GBLUP analysis, therefore 100 stands for a classical numerator relationship matrix (**A**) using pedigree information, 50 is a blended kernel using pedigree and metabolomic information, while 0 represents a relationship matrix created only using the metabolomic data.

Predictive abilities were formally accessed using cross-validation schemes (Figure 5b). As might be expected, results mirrored the variance component analyses (Figure 5a): predictive accuracy increased when more weight is given to metabolites, when compared to pedigree information. The use of kernels based on pedigree records yielded, on average, lower accuracies and large variability across the validation folds. Remarkably, high predictive accuracies and low variation were observed for the ‘Novel Flavor’ when metabolomics data was considered. Combining information from pedigree and biochemical data only leveraged the predictive performance for firmness and texture intensities, both sensory traits associated to physical attributes and easily accessed by instrumental data (i.e., size diameter and compression force).

## 4 Discussion

Over recent decades, it is generally accepted that drop-off in flavor is one of the major sources of consumer dissatisfaction in fruit and vegetable cultivars (Klee, 2010; Klee and Tieman, 2018; Goff and Klee, 2006; Pott *et al*., 2019). With this awareness, modern breeding programs have been seeking for new strategies to make flavor evaluations more efficient and less subjective. Conventionally, flavor acceptability in fruits have been accessed via sensory panels. After pairing to the chemical profile, the results provide a qualitative and quantitative list of potential chemicals associated with the consumer acceptability and can be used to guide breeding selections. Although extensively used, most of the panel surveys in blueberry have overlooked the fact that some genotypes have remarkable aromatic notes that, hypothetically, could impact the consumer experience. However, what makes some genotypes more aromatic than others and what can we learn from that? In this study we dissected the aroma phenotype in a group of tangible metabolites that could be distinguished and were preferred for blueberry consumers. We have unearthed evidence that a group of terpene volatiles enhance consumer acceptability and can be more easily used for flavor prediction.

Identification of metabolites that impact flavor sensation has been an important target to improve the quality of fruits and vegetables (Klee, 2010; Gilbert *et al*., 2014; Farneti *et al*., 2020; Schwieterman *et al*., 2014; Gonda *et al*., 2010; Farcuh *et al*., 2020). Unlike other fruits in which flavor is well-defined by a particular set of orthonasal flavors (e.g., banana and strawberry), the typical blueberry flavor is not controlled by a single chemical component. This fact not only increases the complexity of flavor evaluations, but also inserts a certain subjectivity on the consumer acceptance. A recent metabolomic work carried out in blueberry proved that a large portion of liking perception is dictated by the volatile profile of the fruit (Colantonio *et al*., 2020). Despite the importance, it is still elusive how those volatiles are modulating our flavor inclinations. As a main hypothesis, we leverage our analysis by focusing explicitly on aroma notes, a trait that has been systematically observed by blueberry breeders, but which has not been formally studied and hence had unknown biological mechanisms.

Blueberry aroma was first assayed by classifying a representative group of genotypes regarding the presence of unique aromatic notes and then by pairing sensory and metabolomic analysis. Chemically, over half of the quantity of volatiles collected in both “aromatic” and ‘non-aromatic’ groups were oxylipin derived, confirming earlier literature showing that aldehydes and alcohols make up the majority of blueberry volatiles (Gilbert *et al*., 2015; Farneti *et al*., 2017; Du *et al*., 2011). The main difference between both groups was the quantity of terpenoids. Berries sampled from the “aromatic” genotypes had more than twice the terpene content, when compared to berries from the “non-aromatic” group. But, is this difference noticeable at the sensory level; and, if so, are the “aromatic” berries preferred by consumers? To address this question, we paired chemical information from the fruit with consumer acceptability and observed that panelists systematically rated the “aromatic” berries with higher liking scores. This confirmed our initial hypothesis that such unique flavor notes are preferred by blueberry consumers. Interestingly, “aromatic” berries were also rated as sweeter even though the level of soluble solids and TA – which are standard approximations for berry sugar and acid content – were not found to be statistically different between both groups. It ultimately suggests that some terpene volatiles in blueberries may act by enhancing sweetness, aroma and liking sensation in panelists. Accordingly, similar results endorsing the role of volatiles in enhancing human sweetness perception have been reported in tomato (Tieman *et al*., 2017; Martina *et al*., 2021).

More generally, volatile terpenoids constitute the largest class of plant volatiles and greatly contribute to floral notes emitted by a wide variety of foods (Pott *et al*., 2019). In fruits and vegetables, multiple studies have proved the importance of terpenes for flavor perception. For example, in tomato, over expression of terpenes in genetically modified materials was shown to enhance consumer liking (Davidovich-Rikanati *et al*., 2007). In strawberry, linalool, ethyl butanoate, methyl butanoate, and furaneol; are some terpenes positively associated to consumer liking scores (Schwieterman *et al*., 2014). In carrot, terpene composition represents a critical quality attribute and plays a major role in the production of aroma and flavor compounds (Muchlinski *et al*., 2020). In sweet cherry, on the other hand, more C6 aldehydes and less esters have been suggested to create a herbaceous/grassy-like flavor (Villavicencio *et al*., 2021). In blueberry, although most sensory studies have not explicitly addressed aroma perception in their sensory panels, there is evidence suggesting that linalool, geraniol and nerol contributed to “sweet”, “floral”, “fruity” and “berry-like” aroma notes (Du and Rouseff, 2014). Such aromatic notes agree with a group of lexicons created to described aroma/flavors in blueberry (Bett-Garber and Lea, 2013).

Motivated by the possibility of enhancing fruit flavor, most fruit breeding programs have streamlined flavor evaluations and guide their selections based on sugar and acid concentration. Beside the fact that sugar and acids might not explain the majority of variation associated to flavor acceptability (Colantonio *et al*., 2020), both traits are predominantly controlled by a very tight and multi-layered regulation system (Pott *et al*., 2019). Herein we make the point that aroma – and consequently flavor – can be better accessed via volatile analyses. A primary advantage being that biological pathways associated with secondary metabolites in plants – which include the terpene volatiles – are less complex than sugar and acids and hence more responsive to breeding and selection. For example, several of the terpenes reported here and by other authors have exhibited high levels of heritability and simpler genetic architecture (FerrÃo *et al*., 2020; Bineau *et al*., 2021). Both facts are reinforcing that by selecting for higher or molecularly over-expressing specific terpenes is a more tangible target for fast flavor genetic improvements either via conventional breeding or using molecular tools. Also relevant for breeding, but in a somewhat different perspective, our results are also suggesting that terpene volatiles could be used as a substitute for sugar content by leveraging sweetness perception. Similar results were also reported in tomato, with some volatiles modulating our sweetness acceptability (Martina *et al*., 2021). More broadly, we envision that such findings can significantly impact the human diet, since sweetness perception could be a target without increasing the caloric content of blueberries.

Regarding the use of metabolomics for flavor assisted selection, another recent line of research has suggested the use of all metabolite simultaneously to predict sensory traits – an approach that is analogous to the concept of genomic prediction using DNA information (Meuwissen *et al*., 2001). Discussed in further details by Colantonio *et al*. (2020), we extended the analyses by combining metabolomic and genetic information in the same framework. Our general strategy consisted of predicting consumer acceptability using a variation of mixed model analysis, in which a single blended relationship matrix accounting for pedigree and metabolite information were tested. With that in hand, we could quantify the importance of genetic (accessed via pedigree records) and metabolomic data for flavor predictions. Similar ideas have also been used in other contexts in the plant literature, but not directly applied to fruit flavor improvements. Some recent examples are reported in canola (Knoch *et al*., 2021), oat (Hu *et al*., 2021), rice (Xu *et al*., 2021; Wang *et al*., 2019), wheat (Longin *et al*., 2020), and maize (Schrag *et al*., 2018).

For aroma projection, we observed high predictive accuracy when predictions were performed based on metabolite information. The inclusion of genetic pedigree, in counterpart, did not improve the predictive ability and showed poor results when solely applied. For practical application, we envision that metabolomic analyses could be embraced by other fruit breeding programs. We agree that more cost-effective methods are necessary, particularly because evaluating multiple volatiles simultaneously is still cost prohibitive for breeders (Fernie and Schauer, 2009). Motivated by that, we focused on a selected group of terpenes to explicitly address the aroma classification. We observed that myrtenal, P-cymene and D-limonene were the three volatiles that, individually, led to accurate classifications. Linalool, another terpene commonly reported as a major aroma active volatile in blueberry (Du and Rouseff, 2014) and strawberry (Yan *et al*., 2018), also showed high prediction rates. By focusing on a small subset of key volatiles and developing fast extraction and quantification methods, we envision that blueberry breeding programs could leverage the selection of more flavorful cultivars by increasing the accuracy and reducing time, resources and the high subjectivity associated with flavor evaluations.

More generally, we also projected the combined use of pedigree and metabolomic information for the other characteristics accessed via sensory panel. Similar to the aroma trait, including pedigree information did not improve the predictive ability for most of the traits. In general, we observed mid-to-low predictive performances, a result that has been reported in other crops (Eggink *et al*., 2012). When contrasted to the predictive results previously reported in blueberries by Colantonio *et al*. (2020), herein we noticed lower values – a fact that is probably associated to the smaller training data set used in this study. Nevertheless, in an attempt to confirm the importance of metabolites for flavor prediction, we considered an additional validation step by predicting overall liking in this current data set, but using the regression coefficients originally estimated by Colantonio *et al*. (2020). To this end, we selected all metabolites overlapping both studies, recorded the coefficients estimated for each component using the BayesA model, predicted the overall liking, and correlated the predicted and the real values observed in this present study. Remarkably, a prediction ability larger than 60% was observed. As we argue that this is a true validation exercise – since independent populations were used for calibrations and predictions – we confirm that metabolomic selection can be effectively used for cross predictions and therefore flavor assisted selection.

Finally, we believe that our findings have important applications for blueberry. At the market level, it was previously demonstrated that blueberry consumers are willing to pay more for flavorful foods (Gilbert *et al*., 2014). We believe that this finding opens new opportunities for breeders and growers to distinguish their berries and access premium markets – a strategy effectively used in other fruits (i.e., apple, tomato, and grapes) but not yet well-explored in blueberries.

## 5 Conclusion

In this study we have demonstrated that chemical and sensory data can be combined to support flavor-assisted selection in blueberry. Regarding our main research questions, we can conclude that (i) berries with unique aromatic notes observed in field trials can be perceived and, more importantly, are preferred by blueberry consumers; (ii) a group of terpenes, that include myrtenal, P-cymene, D-limonene and linalool, are key volatiles driving this aroma sensation and hence are potential targets for breeding programs; (iii) with few terpene volatiles we could classify blueberry genotypes as aromatics; and, finally, (iv) the use of metabolomic prediction emerges as a valuable tool to assist with the accurate selection of more flavorfull cultivars and can reduce time, resources and the subjectivity associated with flavor evaluations. Overall, the methods and approaches we used here allow us to provide a blueprint of how blueberry flavor can be understood and opens the frontier to using statistics to predict human flavor interpretation using metabolites. We emphasize that such findings have a broad relevance in the context of plant breeding for fruits. At the economic standpoint, selecting by aromatic fruits may open new opportunities to blueberry growers to distinguish their berries and access a still incipient premium market.

## 6 Acknowledgments

The authors thank David Norden and Werner Collante for their help in the classification of a diverse panel of blueberry genotypes comprising “aromatic” and “non-aromatic” materials; and Mia Acker to help in the data mining. This work was supported by the UF royalty fund generated by the licensing of blueberry cultivars.

## 7 Author Contribution

PRM conceived and supervised the study. HS, PL and PRM classified and created the diversity blueberry panel used in this study. HS and DT performed the volatile extractions, GC-MS data analysis and quality control. CS and HS organized the sensory panels. LFVF and RRA analyzed and interpreted the phenotypic, metabolomic and sensory panel data. LFVF, HS and PRM wrote the paper with revision from all authors. All authors read and approved the final version of the manuscript for publication.

