## Supplementary figures and images for "Terpene volatiles mediates the chemical basis of blueberry aroma and consumer acceptability"

### SM_Fig_1_2_3

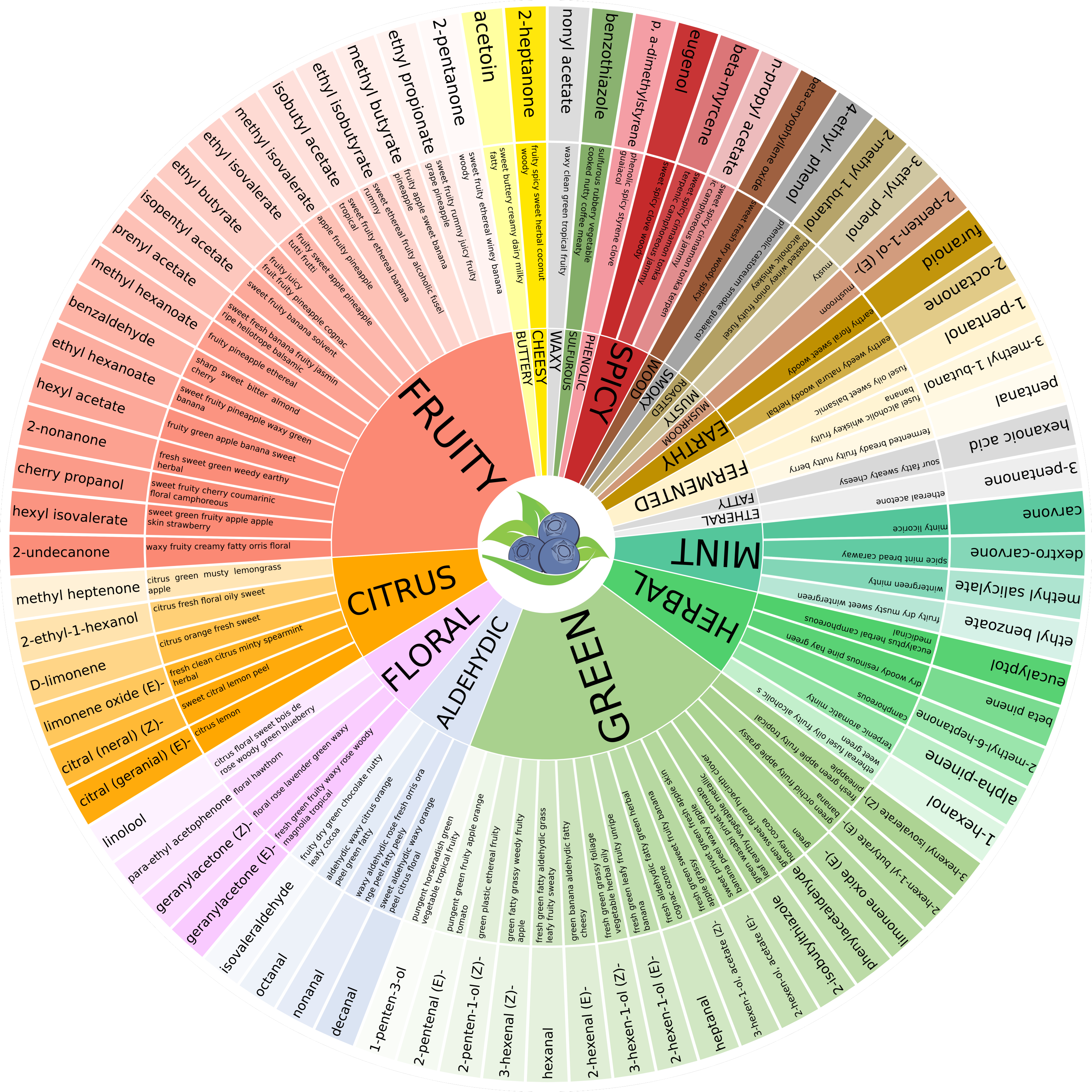

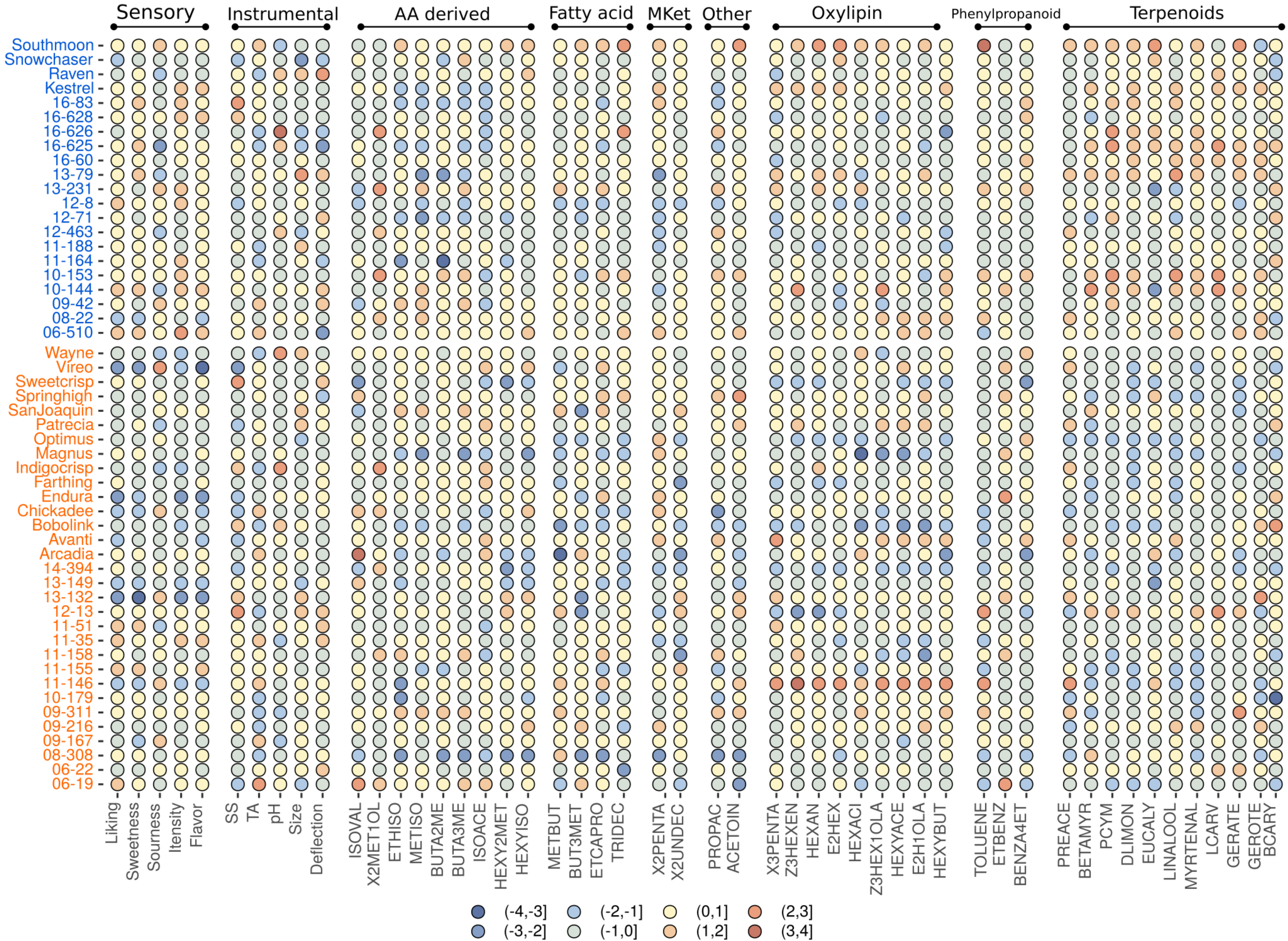

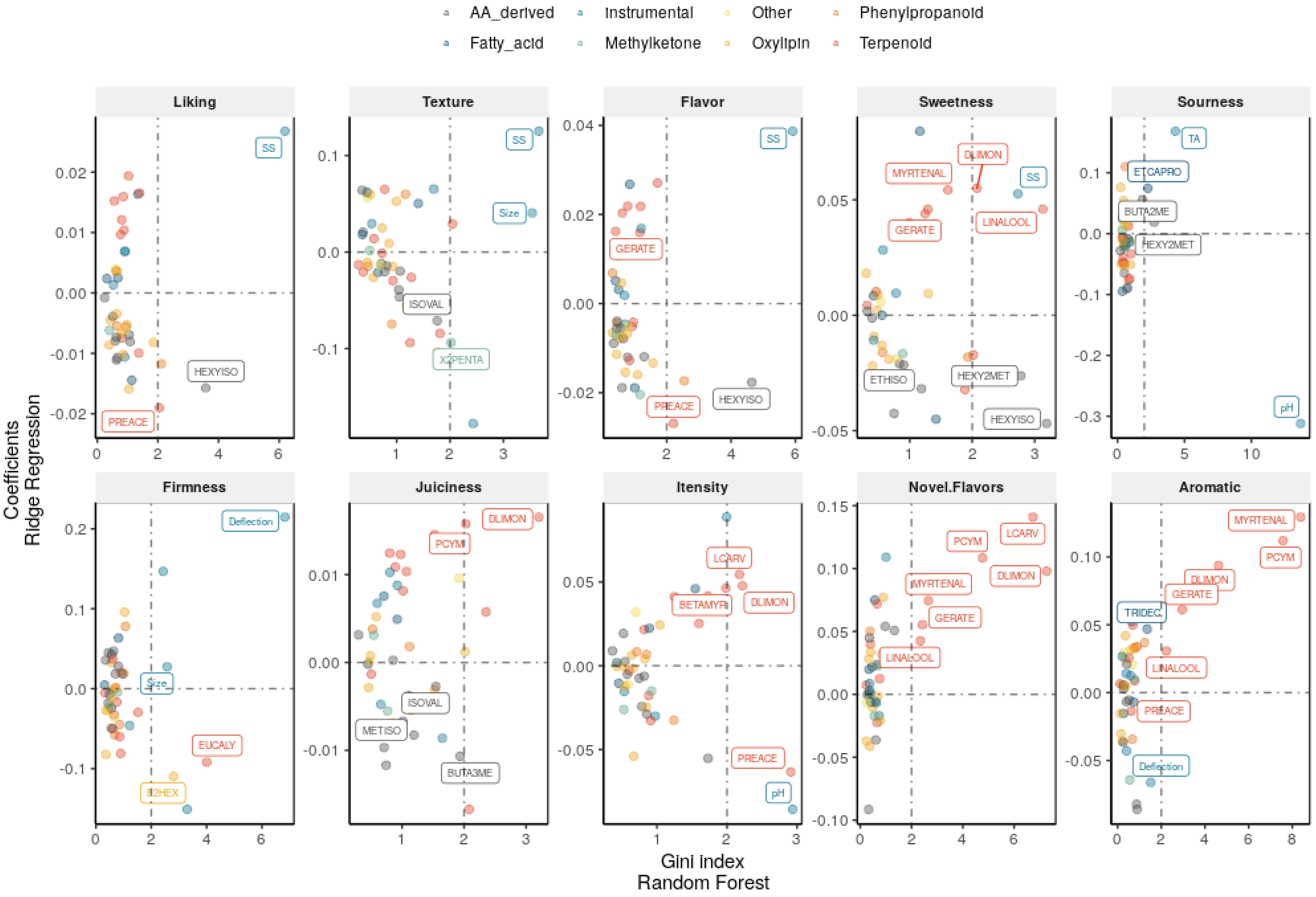
